# Distinct EH domains of the endocytic TPLATE complex confer lipid and protein binding

**DOI:** 10.1101/2020.05.29.122911

**Authors:** Klaas Yperman, Anna C. Papageorgiou, Romain Merceron, Steven De Munck, Yehudi Bloch, Dominique Eeckhout, Pieter Tack, Thomas Evangelidis, Jelle Van Leene, Laszlo Vincze, Peter Vandenabeele, Martin Potocký, Geert De Jaeger, Savvas N. Savvides, Konstantinos Tripsianes, Roman Pleskot, Daniel Van Damme

## Abstract

Clathrin-mediated endocytosis (CME) is the gatekeeper of the plasma membrane. In contrast to animals and yeasts, CME in plants depends on the TPLATE complex (TPC), an evolutionary ancient adaptor complex. The mechanistic contribution of the individual TPC subunits to plant CME remains however elusive. In this study, we used a multidisciplinary approach to elucidate the structural and functional roles of the evolutionary conserved N-terminal Eps15 homology (EH) domains of the TPC subunit AtEH1/Pan1. By integrating high-resolution structural information obtained by X-ray crystallography and NMR spectroscopy with all-atom molecular dynamics simulations, we provide structural insight into the function of both EH domains. Whereas one EH domain binds negatively charged PI(4,5)P_2_ lipids, unbiased peptidome profiling by mass-spectrometry revealed that the other EH domain interacts with the double N-terminal NPF motif of a novel TPC interactor, the integral membrane protein Secretory Carrier Membrane Protein 5 (SCAMP5). Furthermore, we show that AtEH/Pan1 proteins control the internalization of SCAMP5 via this double NPF peptide interaction motif. Collectively, our structural and functional studies reveal distinct but complementary roles of the EH domains of AtEH/Pan1 have in plant CME and connect the internalization of SCAMP5 to the TPLATE complex.

## Introduction

Internalization of membrane proteins is of crucial importance for cell survival as it allows to quickly react to changing environmental conditions. The residence time of integral membrane proteins at the plasma membrane is controlled by internalization signals that are recognized by adaptor protein complexes that mediate a process named clathrin-mediated endocytosis (CME). The start of CME is marked by an enrichment of cargo proteins and negatively charged PI(4,5)P_2_ lipids (Ischebeck et al., 2013). In the next stage, adaptor proteins and clathrin are recruited in a timed fashion and accumulate at the site of endocytosis (Qi et al., 2018). In the later stages, accessory proteins and additional clathrin molecules are recruited followed by scission of the formed vesicle from the plasma membrane. The main driver during this process are low affinity protein-protein and protein-lipids interactions, which enable this dynamic process of assembly and disassembly.

In plants, two adaptor complexes play a role during the initiation phase of endocytosis, the Adaptor Protein-2 complex (AP-2) and the TPLATE complex (TPC) (Gadeyne et al., 2014; Rubbo et al., 2013). AP-2 and TPC likely have independent but also complementary roles in CME (Gadeyne et al, 2014). Both protein complexes have a core-complex of four subunits (Hirst et al., 2014), which in the case of TPC is associated with four additional subunits (TWD40-1, TWD40-2, AtEH1/Pan1 and AtEH2/Pan1). The AtEH/Pan1 proteins are more loosely associated with TPC as they for example do not associate with the other complex subunits when a truncated TML subunit forces TPC into the cytoplasm (Gadeyne et al., 2014) and they do not co-purify with the complex in *Dictyostelium* (Hirst et al., 2014). Recent evidence suggests however a similar arrival time of all TPC subunits at the plasma membrane at the onset of endocytosis, preceding clathrin arrival (Gadeyne et al., 2014; Narasimhan et al., 2020; Wang et al., 2020).

The AtEH/Pan1 proteins are the plant homologs of yeast Pan1p, which is known for its role as an activator of ARP2/3-dependent actin dynamics during endocytosis (Duncan et al., 2001; Toshima et al., 2005, 2007). Pan1p was also recently shown to be part of a phosphorylation-dependent mechanism connecting endocytic vesicles, endosomal compartments and actin dynamics in budding yeast (Toshima et al., 2016). The Arabidopsis AtEH/Pan1 proteins were recently shown to mediate actin-dependent autophagy in plants (Wang et al., 2019). AtEH/Pan1 and Pan1p proteins are both hallmarked by the presence of two EH domains at their N-terminus (Gadeyne et al., 2014; Wang et al., 2019).

In animals and yeast, EH domains have been characterized in great detail due to their presence in crucial endocytic proteins like Eps15, REPS1, EHD1, etc. (Beer et al., 2000; Kieken et al., 2009; Kim et al., 2001). SMART and Prosite analysis identified only six EH domains in Arabidopsis compared to eighteen in humans (Letunic and Bork, 2017; Sigrist et al., 2013). In Arabidopsis, EH domains are present in the endocytic recycling regulators EHD1 and EHD2, each having one EH domain (Bar et al., 2008) and in the TPC subunits AtEH1/Pan1 and AtEH2/Pan1, each having two EH domains. Plant EHD1 and EHD2 proteins have been characterized as homologues of human EHD proteins and were suggested to play a regulatory role during plant CME, plant defense and salt stress (Bar and Avni, 2009; Bar et al., 2008, 2013). In contrast to the essential function of all tested TPC subunits, silencing of EHD1/2 does not result in severely aberrant phenotypes, indicating redundancy or a more specialized function (Bar et al., 2008; Gadeyne et al., 2014; Wang et al., 2019). In contrast to the single EH domain in EHD proteins, AtEH/Pan1 proteins harbor two EH domains and we asked if both EH domains serve as independent functional modules. We therefore set out to characterize the function of the AtEH/Pan1 EH domain containing proteins in Arabidopsis. In this study, we used a multidisciplinary approach to perform a structural and functional side-by-side comparison of both EH domains of AtEH1/Pan1.

## Results and Discussion

### Structural characterization of both EH domains of AtEH1/Pan1 reveals a common fold

AtEH/Pan1 proteins are highly unstructured but three domains can be identified; a coiled-coil domain implicated in dimerization (Sánchez-Rodríguez et al., 2018) and two N-terminal Eps15 homology (EH) domains (Figure 1, panel a). Comparing the EH domains within each AtEH/Pan1 protein shows low sequence identity which is in contrast to the high sequence identity when comparing the EH domains between AtEH1/Pan1 and AtEH2/Pan1 (Figure 1, panel a; Supplementary Figure 1). Therefore, we decided to focus on both EH domains of AtEH1/Pan1 as representatives, which we hereafter name EH1.1 and EH1.2. To structurally characterize both EH domains of AtEH1/Pan1, we expressed recombinant proteins in *E.coli* and purified highly monodisperse samples for X-ray crystallography and NMR (Supplementary Figure 1). Crystals of EH1.1 diffracted synchrotron X-rays to 1.55Å resolution and enabled structure determination via molecular replacement (PDB:6YIG). EH1.1 consists of two EF-hands connected by a short antiparallel β-sheet (Figure 1, panel b). Anomalous scattering supports a calcium ion bound in a pentagonal-bipyramidal geometry in the loop of the first EF-hand (Figure 1, panel d; Supplementary figure 1). No anomalous scattering signal was detected in the second EF-hand loop, but electron density consistent with the coordination of a sodium ion was present (Figure 1, panel c). While attempts to crystallize EH1.2 proved unsuccessful, and as part of an integrative structural biology approach, we obtained structural insights for both EH1.1 (PDB: 6YEU) and EH1.2 (PDB: 6YET), in solution, by NMR (Evangelidis et al., 2018). With respect to EH1.2, the NMR and X-ray structures agree very well with an RMSD of 1Å (compared by the Dali algorithm ((Holm, 2019)). When comparing the NMR structures of EH1.1 and EH1.2, the RMSD between both domains is 2.6Å (compared by the Dali algorithm ((Holm, 2019)). In general, the EH1.1 and EH1.2 domains have a very conserved hydrophobic core, a common feature of EH domains. The major difference in the fold is at the interaction interface of the N- and C-terminal ends. The hydrophobic core in EH1.1 is shielded by the proline-rich C-terminal loop, while in EH1.2, the proline-rich N-terminal loop takes over this function.

**Figure 1:**
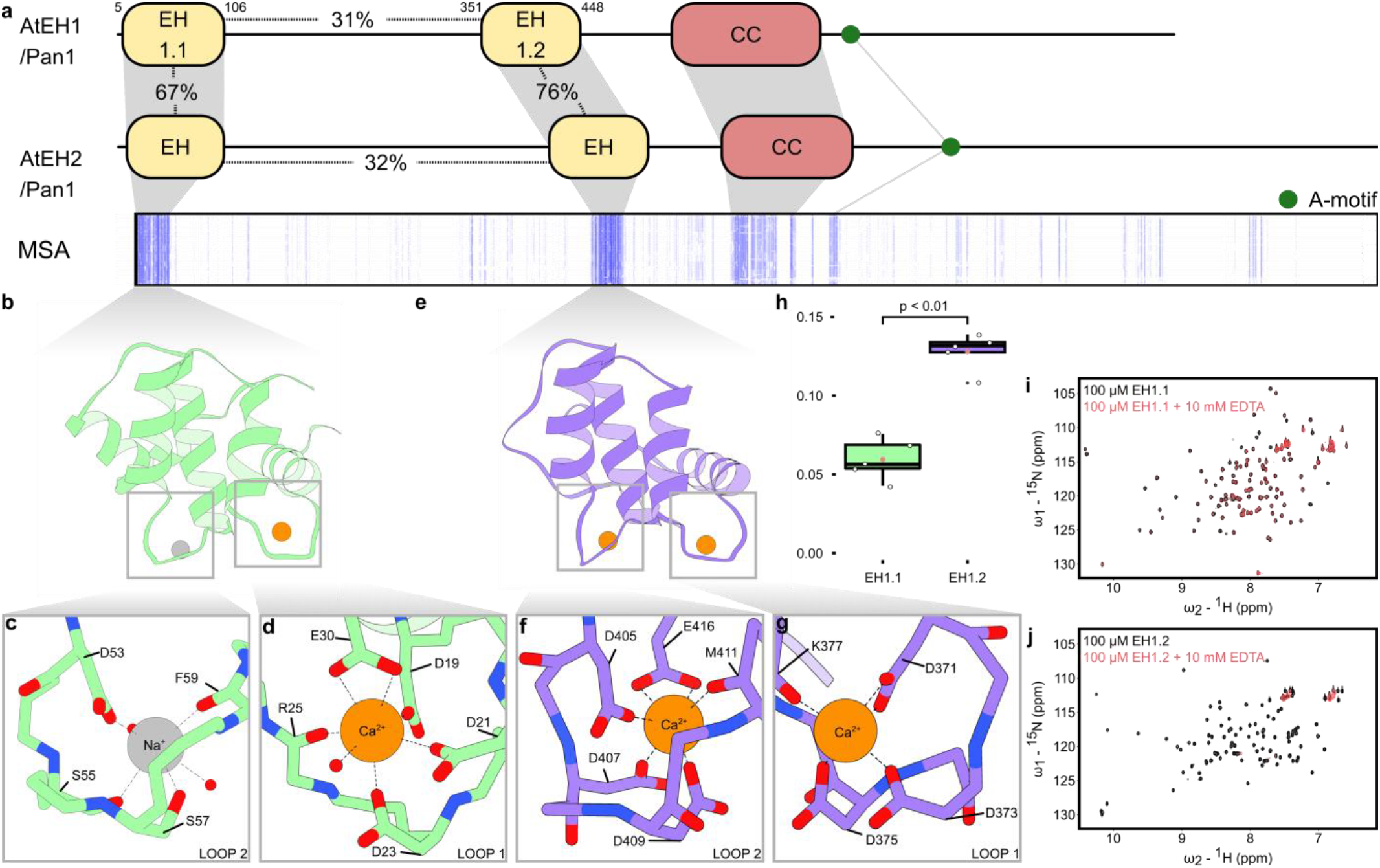
EH domains of AtEH1/Pan1 differ in their Ca^2+^-binding capacities. **a**, Domain organization of AtEH1/Pan1 and AtEH2/Pan1. Both proteins contain two Eps15 homology domains (EH), a coiled-coil domain (CC) and an acidic (A)-motif. A schematic representation of a multiple sequence alignment (MSA), shows strong conservation of the EH domains (blue lines) across the plant kingdom. Percentages indicate the relative number of identical amino acids. **b-g**, Cartoon representation of the X-ray structure of EH1.1 and NMR/all-atom molecular dynamics structure of EH1.2. Ions are shown as orange (Ca^2+^) or grey (Na+) spheres. Insets show the ion coordination in each EF-hand loop. **h**, Total X-ray reflection fluorescence intensities of Ca^2+^ normalised to Cl^-^ of samples containing 2mM of each EH domain in the presence of 0.5mM free Ca^2+^. **i-j**, Superimposed ^15^N-^1^H HSQC spectra of the EH domains before (grey) and after (red) Ca^2+^ chelation by 10mM EDTA.

### The second EH domain coordinates two calcium ions

We were intrigued by the fact that, in contrast to EH1.1, EH1.2 contains two possible calcium-binding sites. This is manifested by a classical calcium-binding motif (DxDxDxxxxxE) in the second loop of EH1.2 and a second possible coordination motif in the first loop, where the final glutamate, in the canonical calcium-ligation cassette, is substituted by glutamine in arabidopsis. However, sequence alignment (supplemental data) revealed that most monocots have glutamate at this position, strongly suggesting calcium coordination in this position. To obtain a possible coordination scheme for arabidopsis we performed an extensive all-atom molecular dynamics simulation (3 μs, CHARMM force-field) (Figure 1, panel e-g, Supplementary Figure 2). Experimental interrogation of this possibility by total X-ray reflection fluorescence confirmed the presence of two calcium ions in the EH1.2 domain (Figure 1, panel h). To our knowledge, the ability of EH domains to coordinate two calcium ions has not been described before. The all-atom molecular dynamics model, within its time and forcefield limitations, suggests that the first aspartate and the presence of an extra water molecule compensate for the incomplete calcium-binding motif in the first loop of EH1.2 and functionally mimics the role of the canonical glutamate in the calcium-binding motif. Restoring the first EF-hand loop to a canonical EF-hand (Q382E), in an all-atom molecular dynamics simulation, resulted in a classical arrangement where glutamate coordinates calcium in a bidentate fashion (Supplementary figure 2). We confirmed by NMR and size exclusion chromatography that EH1.2 is indeed more sensitive to precipitation upon calcium chelation compared to EH1.1 (Figure 1, panel i-j and Supplemental Figure 3). This is consistent with our findings that EH.2 coordinates two calcium ions. Addition of an excess amount of calcium allowed refolding of the precipitated domains (Supplemental Figure 3). The ability to unfold and refold, relating to a non-functional versus a functional state, in a calcium-dependent manner, hints at a modulatory role for calcium to control the function of this domain.

### The second EH domain interacts with charged lipids

EH domains have been proposed to act as a protein interaction hub and/or as a lipid-binding module (Naslavsky et al., 2007; Paoluzi et al., 1998). To unravel the function of the EH domains of AtEH1/Pan1, we tested both domains for their ability to bind peptide motifs and membranes, in a pairwise manner. AtEH/Pan1 proteins mainly function at the negatively charged plasma membrane or the ER-PM contact sites (Gadeyne et al., 2014; Wang et al., 2019). To mimic a negatively charged plasma membrane environment *in vitro* we performed liposome binding experiments using a mixture of phosphatidylcholine (PC) with or without 10% PI(4,5)P_2_. Only EH1.2 bound to PI(4,5)P_2_ enriched liposomes (Figure 2, panel a). We hypothesized that evolutionary conserved lysine and arginine residues at the surface of EH1.2 would result in an electrostatically driven membrane interaction (Supplementary Figure 4). To test our hypothesis we performed liposome binding experiments comparing PI(4,5)P_2_ with PI3P and PI4P enriched liposomes. We observed a very weak interaction with PI3P and PI4P compared to PI(4,5)P_2_ liposomes (Figure 2, panel b). PolyPiPosome assays confirmed our findings (Supplementary Figure 4), along with conservation of specific lysine and arginine residues in the EH domains of Eps15 (EH2) and EHD1 (EH1), both known for their involvement in binding negatively charged lipid head groups (Naslavsky et al., 2007) (Figure 2, panel c). The lipid interacting residues in the EH domains of EHD1 and Eps15 are structurally conserved in EH1.2 (K391 and K398), whereas no lysine or arginine residues are present at those positions in EH1.1 (Figure 2, panel d). In addition to the published interacting residues, we hypothesize a third residue, K384, might also play a role in lipid interaction as it is located close to the known interacting surface and is structurally conserved in lipid binding EH domains (Figure 2, panel c). Our analysis supports that the EH1.2 interaction with PI(4,5)P_2_ liposomes is electrostatically driven and might be *in vivo* further enhanced via avidity effects through dimerization of the AtEH1/Pan1 coiled-coil domain (Sánchez-Rodríguez et al., 2018). Given the fact that, on the one hand, calcium is needed for the fold of EH1.2 and, on the other hand, that the lipid interaction is electrostatically-driven, we tested the effect of different calcium concentrations on PI(4,5)P_2_ binding by the EH1.2 domain. We observed increased binding of EH1.2 at lower calcium concentrations (Figure 2, panel e). Our results are in line with prior studies showing that high calcium concentrations block the accessibility of charged lipids by shielding and re-arranging the lipid headgroups (Bilkova et al., 2017).

**Figure 2:**
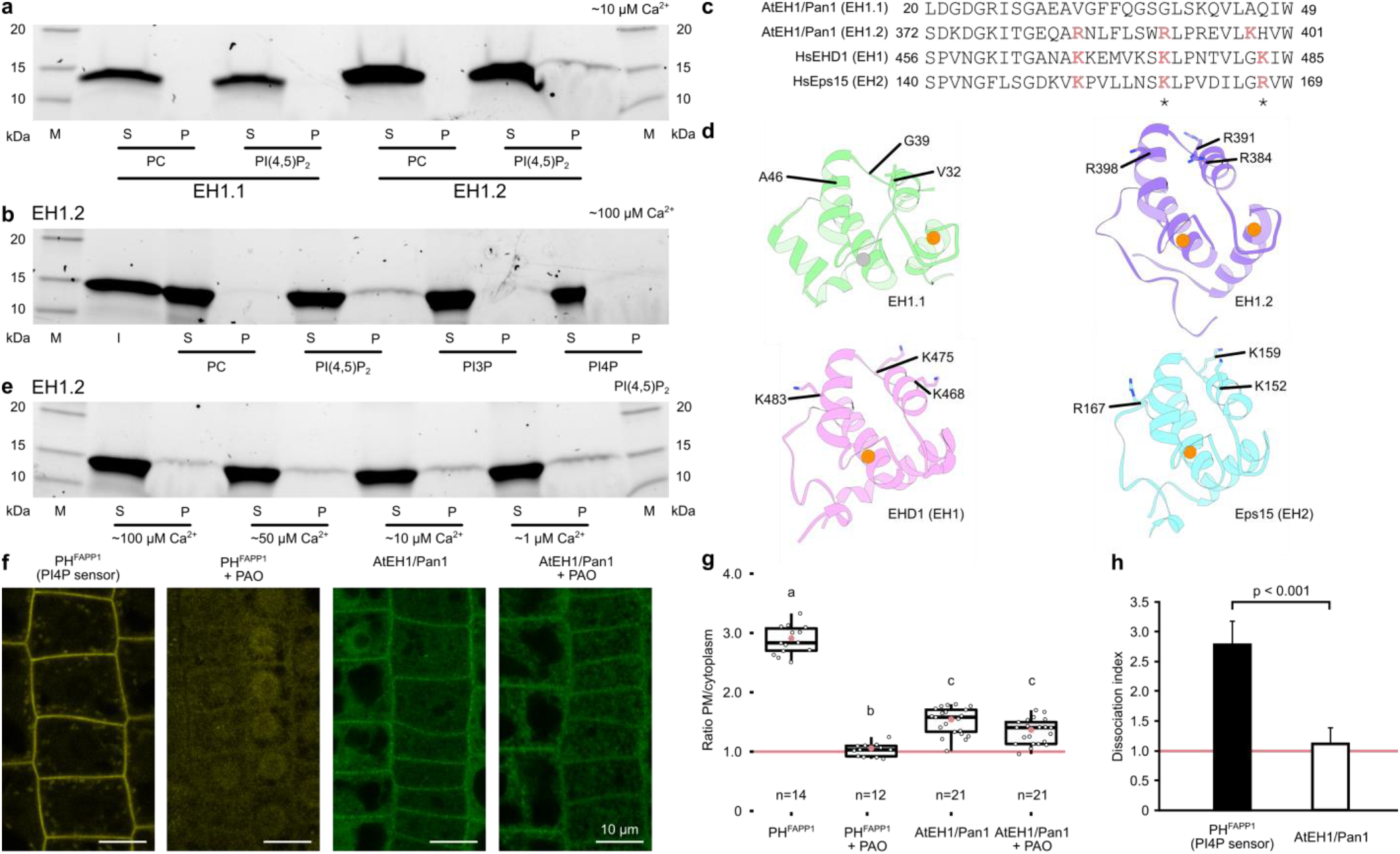
The second EH domain of AtEH1/Pan1 exhibits a Ca^2+^ dependent, charge-based PI(4,5)P_2_ lipid binding. **a**, Coomassie-blue stained SDS-PAGE (4-20%) analysis of liposome binding comparing the binding of the EH domains between PC and 10% PI(4,5)P_2_ containing liposomes. M=marker, S=soluble fraction, P=pellet. **b**, Liposome binding assay of EH1.2 between differently charged liposomes. **c**, Sequence alignment of AtEH1/Pan1 in comparison to EH domains shown by NMR to bind lipids. Conserved lysine and arginine residues are highlighted in pink. Residues shown to bind lipids by NMR are indicated with an asterisk (Naslavsky et al., 2007). **d**, Mapped residues highlighted in panel c indicated on a cartoon representation of EH1.1, EH.2, EHD1 (2KSP) and Eps15 (1F8H). **e**, Liposome binding assay of EH1.2 in a buffer containing different Ca^2+^ concentrations. **f**, Arabidopsis transgenic lines overexpressing CITRINE-PH^FAPP1^ or AtEH1/Pan1-GFP were treated with 30 μM PAO for 30min. AtEH1/Pan1 retained its plasma membrane localization in contrast to the PI4P marker. A representative image for both used lines is shown before and after treatment. Scale bar indicates 10 μm. **g**, Quantification of PAO treatment by plasma membrane versus cytoplasm of three plants. The total number of cells quantified is indicated below each graph. The red line indicates an equal intensity between the plasma membrane and the cytoplasm. Different letters indicate significant differences between samples by Tukey multiple pairwise-comparisons (P < 0.001). **h**, The corresponding dissociation index of panel g.

To address the lipid-binding capacity *in planta*, we assessed if the targetting of AtEH1/Pan1 to the plasma membrane depends on PI4P binding, as no potent compound exists to disrupt PI(4,5)P_2_ concentrations *in vivo.* We used short term (30min), 30μM phenyl arsine oxide (PAO) treatment, shown to specifically affect PI4P levels at the PM (Simon et al., 2014). In contrast to the PI4P marker (PH_FAPP1_-GFP), PAO treatment did not disrupt the AtEH1/Pan1 PM localization indicating that PI4P is not of major importance for the plasma membrane targeting of AtEH1/Pan1 (Figure 2, panel f-h).

### The first EH domain recognises a novel retrograde transport motif

To identify protein interaction partners of the AtEH1/Pan1 EH domains, we sought to discover interaction motifs. To this end, we digested arabidopsis seedling proteome and incubated the peptide mix with each EH domain or with GFP as a control. Comparative mass spectrometry identified bound peptides. The N-terminal double NPF peptide of SCAMP5 was identified as a significant hit with EH1.1. This peptide also showed the hightes fold change compared to EH1.2 and GFP (Figure 3, panel a). Secretory Carrier Membrane Protein 5 (SCAMP5) is part of a five-membered protein family in arabidopsis. SCAMP proteins were first characterized in mammals for their role in endocytosis (Fernández-Chacón et al., 2000). Current insights in the animal and plant field show a broader function of SCAMPs in plasma membrane phase separation, cell plate formation, and pathogen-induced stomatal closure (Bourdais et al., 2019; Lam et al., 2008; Park et al., 2018). SCAMP5 was also previously identified to reside in close proximity to TPC by proximity labelling (Arora et al., 2020). No peptides derived from integral membrane proteins were identified among the few specific interactors of EH1.2 (Bateman et al., 2018; Letunic and Bork, 2017).

**Figure 3:**
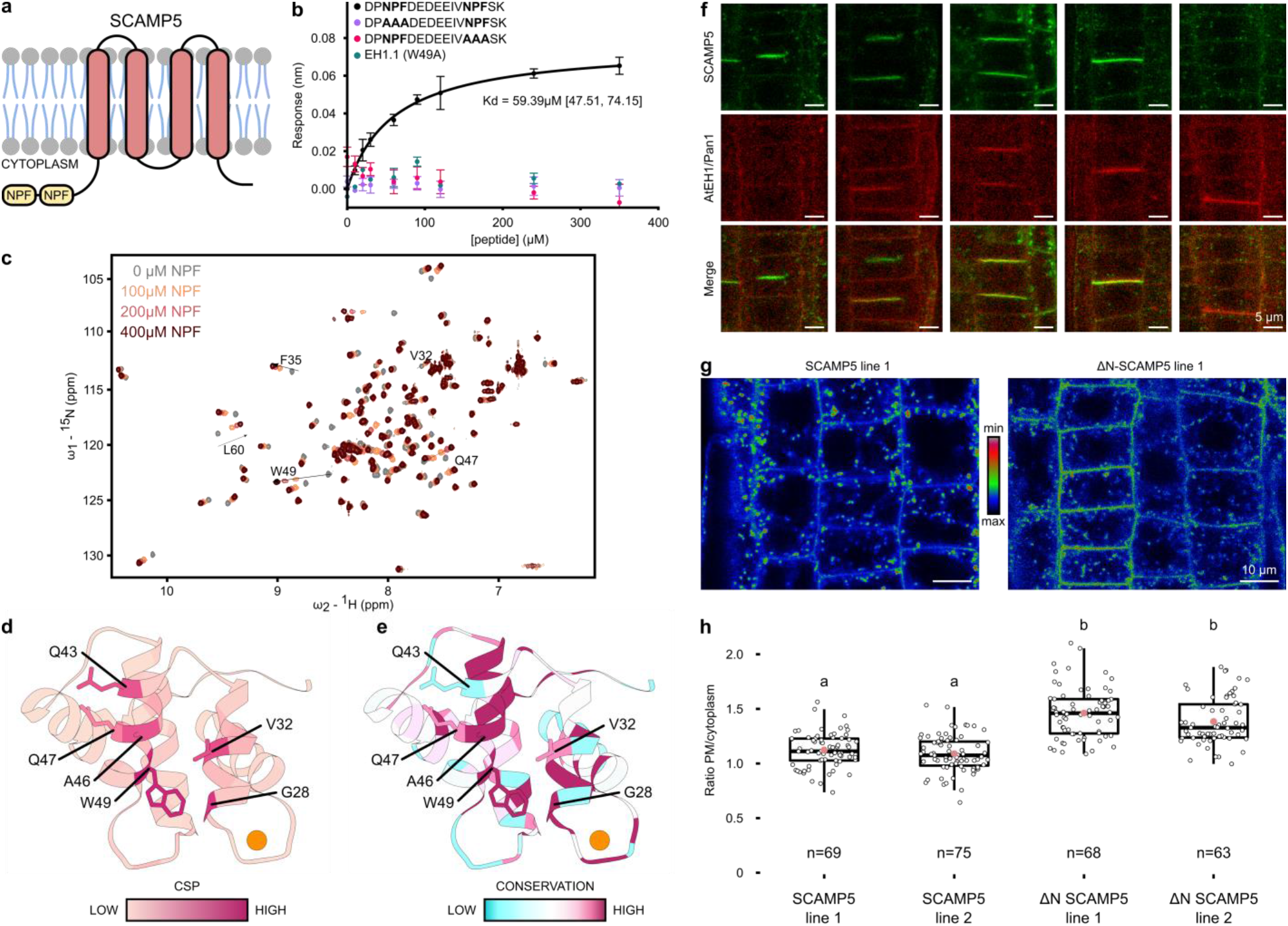
The first EH domain of AtEH1/Pan1 interacts with the N-terminal double NPF motif of SCAMP5. **a**, Graphical representation of SCAMP5, the double N-terminal NPF motif is indicated in yellow. **b**, BLI steady-state kinetics of the binding of EH1.1 of AtEH1/Pan1 and NPF peptides with or without mutations. WT Protein but not the W49A mutant binds the double NPF peptide with a measurable affinity. Mutation of any of the NPF motifs abrogates binding. **c**, ^1^H-^15^N-HSQC spectra of EH1.1 titrated with increasing amounts of the SCAMP5 double NPF peptide (grey to red). Peak trajectories of selected residues are indicated by arrows. Their weighted chemical shift perturbations were used to obtain binding isotherms and derive an apparent dissociation constant of the interaction (Supplementary Figure 6). **d**, Cartoon representation of EH1.1 (6-105) colored with a gradient (pink to purple) indicate the extent of chemical shift perturbations induced by the SCAMP5 double NPF peptide binding. Residues showing large chemical shift perturbations (> 0.3ppm) are shown as sticks. **e**, Similar as in d but the EH1.1 structure is colored according to ConSurf colors denoting evolutionary conservation (blue to white to purple). Most residues affected by peptide binding are well-conserved. **f**, Colocalization of SCAMP5-GFP and AtEH1/Pan1-mRuby3 at the plasma membrane and the cell plate in Arabidopsis root cells. Cells in different phases of cytokinesis are depicted. SCAMP5 recruitment to the cell plate precedes AtEH1/Pan1 whereas the presence of the latter at the newly formed cross wall exceeds SCAMP5 following completion of cytokinesis. Scale bar indicates 5 μm. **g**, Confocal analysis of SCAMP5-GFP vs ΔN-SCAMP5-GFP. An increased plasma membrane localization was observed in the absence of the double NPF motif. Colors are shown according to signal intensity (red to green to blue). Scale bar indicates 10 μm. **h**, Quantification of the plasma membrane vs the cytoplasm of two independent lines for both constructs as shown in panel g. The mean is shown as a pink dot. The amount of quantified cells is indicated below each boxplot. Different letters indicate significant differences between samples by Tukey multiple pairwise-comparisons (P < 0.001).

Bio-Layer Interferometry (BLI) confirmed the interaction between EH1.1 and a peptide derived from the N-terminus of SCAMP5 with a binding affinity (K_d_) of 59μM via steady-state kinetics (Figure 3, panel b). Both NPF stretches of the N-terminus of SCAMP5 are required to bind EH1.1, as no binding affinity for mutated NPF to AAA mutants could be determined (Figure 3, panel b). The interaction between the double NPF motif is much stronger for EH1.1 (K_d_,^app^ ~ 33μM) compared to EH1.2 (K_d_,^app^ ~190 μM) by NMR titration experiments (Supplementary Figure 6). NMR peptide titration experiments provided the binding sites of the peptide on both EH domains. Published structures of EH domains interacting with NPF-motifs show the importance of the central tryptophan and its surrounding hydrophobic residues for which in EH1.1, but not EH1.2, the largest chemical shift perturbations were observed (Beer et al., 2000; Kieken et al., 2009; Rumpf et al., 2008) (Figure 3, panel c-d, Supplementary Figure 6). Comparison of chemical shift perturbation of EH1.1 with evolutionary conservation across a variety of plant species showed that most of the residues responsible for NPF binding are strongly conserved (Figure 3, panel e). Mutating the conserved tryptophan to alanine in EH1.1 and testing its binding capacity by BLI confirmed its essential role. No binding was observed for the mutated EH1.1 domain (Figure 3, panel b). We conclude that the N-terminal double NPF motif of SCAMP5 interacts specifically with EH1.1 via a hydrophobic interaction mediated by the conserved tryptophan residue.

To address the physiological relevance of the interaction between SCAMP5 and AtEH1/Pan1 *in planta*, we analyzed *A.thaliana* roots expressing SCAMP5-GFP and AtEH1/Pan1-mRuby3 via confocal microscopy. SCAMP5 localizes mostly in endosomes and weakly at the plasma membrane (Supplemental figure 7). Co-localization with AtEH1/Pan1 at the PM was observed. Both proteins also prominently co-localize during various stages of cell plate formation where SCAMP5 clearly precedes the arrival of AtEH1/Pan1. However, the presence of AtEH1/Pan1 at the newly formed cross wall exceeds SCAMP5 following completion of cytokinesis (Figure 3, panel f). Altogether our data suggest that SCAMP5 trafficking is highly dynamic. Short-term ES-9 treatment, a potent endocytic inhibitor (Dejonghe et al., 2016), caused SCAMP5 accumulation at the PM (Supplemental Figure 7), indicating the endocytic contribution to SCAMP5 dynamics. To further elucidate the role of the AtEH1/Pan1-SCAMP5 interaction *in vivo*, we compared the localization of the native SCAMP5 protein with an N-terminally truncated version (i.e. lacking the double NPF motif). In comparison to the wild type protein, the ΔN-SCAMP5 showed a reduced endosomal and an increased plasma membrane localization (Figure 3, panel g-h). Altogether, these results corroborate our hypothesis that the double NPF motif is a recruitment signal that is involved in the retrograde transport of SCAMP5.

In conclusion, our parallel structural and functional comparison of the EH domains of AtEH/Pan1 revealed two divergent EH domains with differential, yet complementary functions. The first EH domain binds a novel TPC interactor, SCAMP5, via its N-terminal double NPF motif. This constitutes a novel retrograde transport signal in plants. The second EH domain is involved in the recognition of negatively charged phospholipids. The tandem EH domains in AtEH1/Pan1, a recurring leitmotif in the endocytic machinery in Eukarya, is a clear example of evolutionary division of labour of a repetitive protein fold.

## Acknowledgements

The authors would like to thank Prof. Eugenia Russinova (Ghent University/VIB-PSB) for an aliquot of the ES-9 compound. Prof. Ray Owens (OPPF, Research Complex, Harwell) for initial work on the EH constructs (PID:1724/1846) and an aliquot of the GFP-his plasmid.

This research in the D.V.D lab is supported by the European Research Council T-Rex project number 682436 and by the National Science Foundation Flanders (FWO; G009415N). The research in the K.T lab is supported by project CEITEC 2020 (no. LQ1601) with financial contribution from the MEYS CR and National Programme for Sustainability II and by Grant Agency of Masaryk University (MUNI/G/0739/2017). CIISB research infrastructure project LM2018127 funded by MEYS CR is gratefully acknowledged for the financial support of the measurements at CEITEC Josef Dadok National NMR Centre. We thank the staff of beamlines P14 (PETRAIII) and Proxima2A (SOLEIL) for beam time allocation and excellent technical support. Y.B. was supported by a post-doctoral research fellowship from Research Foundation Flanders (FWO, Belgium). S.D.M. was supported by a pre-doctoral fellowship from the Flanders Agency for Innovation and Entrepreneurship (VLAIO-Flanders, Belgium). S.N.S. acknowledges research support from the Hercules Foundation (no. AUGE-11-029), Ghent University (BOF17-GOA-028) and the VIB. We would like to thank iNEXT (PID: 6554) for funding the structure determination of EH1.1.

## Author contributions

K.Y and R.P designed and performed most of the experiments and wrote the paper with D.V.D. A.P. performed NMR structure calculations calcium precipitation experiments (NMR) and peptide titration experiments (NMR); T.E. helped with structure calculations; R.M. performed MALS and X-ray structure determination and helped in designing EH constructs and protein cystallization; S.D.M. performed BLI; Y.B collected X-ray data and assisted in structure determination; D.E performed MS analysis; M.P helped in performing liposome binding asssays; P.T. performed TXRF.

K.T., S.N.S., G.D.J., L.V., J.V.L., R.P. and D.V.D. were responsible for experimental design, research supervision and finalizing the manuscript text.

## Materials and methods

### Multiple sequence alignment

To obtain protein sequences of AtEH/Pan1 homologs, GenBank (https://www.ncbi.nlm.nih.gov/genbank/), Joint Genome Institute (https://genome.jgi.doe.gov/portal/), EnsemblPlants (https://plants.ensembl.org/index.html) and Congenie (http://congenie.org/start) databases were used for a BLASTP search (Altschul et al., 1990). See supplemental sequence file for a complete list of all organisms searched (51 different plant genomes in total). A multiple alignment was constructed with the masfft algorithm in einsi mode (Katoh et al., 2017).

### Protein production and purification

EH domains of AtEH1/Pan1 were amplified from the pDONR plasmid containing the full-length AtEH1/Pan1 coding sequence (Gadeyne et al., 2014) and cloned by restriction digestion (Nde/Xho) into the pET22b plasmid, generously donated by the lab of S.N. Savvides (IRC, VIB/UGent, BE). The final constructs have an N-terminal His-tag followed by a TEV-protease cleavage site and contain amino acids, 1-107 (EH1.1) and 346-449 (EH1.2). To generate the tryptophan mutant in EH1.1, the complete plasmid was amplified using primers over the tryptophan-containing sequence. The fragment was re-assembled using NEBuilder^®^ HiFi DNA Assembly Master Mix (NEB). Constructs were transformed into BL21(DE3) (#C2527H, NEB). Cells were grown at 37°C in LB^+^ medium and induced by the addition of 0.4mM IPTG at OD 0.6 for 5h. To obtain isotope labelled proteins, cells were grown in M9 minimal medium supplemented with 0.5g/L N15H4Cl and/or 2g/L ^13^C-glucose (Eurisotop). Proteins were extracted using sonication in 20mM HEPES pH 7.4, 150mM NaCl, 2mM CaCl_2_ and Protease inhibitors (cOmplete ULTRA EDTA-free, Roche), except for proteins analyzed by the size exclusion chromatography multi-angle laser light scattering, for which no CaCl_2_ was added during the purification. Purification was performed on an ÄKTA (GE Healthcare) system by subsequent purification using a HisPrep FF 16/10 (GE Healthcare) followed by a gel filtration step (HiLoad^®^ 16/600 Superdex^®^ 75 pg (GE Healthcare)). When no His-tag was required the protein was incubated overnight with 1/40 protein: his-TEV-protease (own production, the expression strain was a gift from the lab of S. Savvides (IRC, VIB/Ugent, BE)) at room temperature without shaking. Uncleaved protein and protease were removed via reverse IMAC (1ml HisTRAP FF (GE Healthcare) followed by gel filtration (HiLoad^®^ 16/600 Superdex^®^ 75 pg (GE Healthcare)). The protein sequence of EH1.1 and EH1.2 along with there native molecular weight were verified by MS analysis.

GFP-His in an OPINF backbone (a generous gift from the lab of Ray Owen, OPPF, UK) was produced and purified as the EH domains without the addition of CaCl_2_.

### Multi-Angle Laser Light Scattering

Purified His-tagged proteins EH1.1 (1mg/ml) or EH1.2 (2mg/ml) were injected onto a Superdex 75 Increase 10/300 GL size exclusion column (GE Healthcare), equilibrated with 20mM HEPES pH 7.4, 300mM NaCl, coupled to an online UV-detector (Shimadzu), a mini DAWN TREOS (Wyatt) multi-angle laser light scattering detector and an Optilab T-rEX refractometer (Wyatt) at room temperature. A refractive index increment (dn/dc) value of 0.185 ml/g was used. Band broadening corrections were applied using parameters derived from BSA injected under identical running conditions. Data analysis was carried out using the ASTRA6.1 software.

### Protein crystallization of EH1.1

Commercial sparse matrix sitting drop crystallization screens were set up using a Mosquito liquid handling robot (TTP Labtech) using a 100nl:100nl, protein (12mg/ml): mother liquor geometry in SwissSci 96-well triple drop plates. Plates were incubated at 293 K. An original hit in the JCSG screen (1.1M SodiumMalonate, 0.1M HEPES pH 7, 0.5% Jeffamine) was optimized to 100mM HEPES pH 7.6, 0.8M SodiumMalonate, 0.5% Jeffamine. Crystals were cryoprotected by the addition of ethylene glycol (15% v/v) to the mother liquor prior to plunging the crystals in liquid nitrogen for cryo-cooling prior to data collection.

### Crystallographic structure determination

X-ray diffraction data were collected from single crystals at 100 K at the P14 microfocus beamline operated by the EMBL at PETRA III synchrotron (Hamburg, Germany). All data were integrated and scaled using XDS (Kabsch, 2010). The initial phases were generated by Automatic Molecular Replacement Pipeline (MoRDa) (Vagin and Lebedev, 2015) using a search model derived from the X-ray structure of mouse EHD2 (2QPT). The initial structure was rebuilt with ARP/wARP (Langer et al., 2008) and further structure building and refinement was performed using Buster (version 2.10.3) (Bricogne et al., 2018) followed by iterative use of COOT (Emsley et al., 2010) and Phenix.refine (Adams et al., 2010) software packages.

### NMR structure determination of EH1.1 and EH1.2

For NMR structure determination the proteins were buffer exchanged using PD-10 columns (Sephadex G-25 M, GE Healthcare) or via gel filtration chromatography in the final purification step to 20mM MES 6.5, 150mM NaCl, 2mM CaCl_2_. All NMR spectra were recorded at CEITEC Josef Dadok National NMR Centre on 850 MHz Bruker Avance III spectrometer equipped with 1H/13C/15N TCI cryogenic probehead with z-axis gradients. For each protein a set of three sparsely sampled 4D NMR experiments was acquired: 4D HC(CC-TOCSY(CO))NH, 4D ^13^C,^15^N edited HMQC-NOESY-HSQC (HCNH), and 4D ^13^C,^13^C edited HMQC-NOESY-HSQC (HCCH). Sequential and aliphatic side chain assignments were obtained automatically using the 4D-CHAINS algorithm that combines through-bond information from the 4D-TOCSY experiment and distance information from the 4D-NOESY (HCNH) experiment (Evangelidis et al., 2018). Aromatic sidechain frequencies were assigned manually by recording an additional 3D ^13^C edited NOESY-HSQC experiment. Assignment completeness reached 99% for each EH domain. Backbone dihedral angle restraints were derived from TALOS using a combination of five kinds (H^N^, H^α^, C^α^, C^β^, N) of chemical shift assignments for each residue in the sequence (Shen et al., 2009). NOE cross-peaks from the three NOESY spectra were assigned automatically by CYANA 3.0 in structure calculations with torsion angle dynamics (Güntert, 2008). Unambiguous distance restraints and torsion angle restraints (Supplementary Table 2) were used in a water refinement calculation (Linge et al., 2003) applying the RECOORD protocol (Nederveen et al., 2005). The CNS patch introducing calcium coordination in a pentagonal bipyramidal configuration was prepared manually. CNS topology files for calcium coordination were generated based on the high-resolution crystal structure of calmodulin (PDB:1CLL). The quality of the NMR-derived structure ensembles was validated using PSVS (Bhattacharya et al., 2006).

### All-atom molecular dynamics simulations of Ca^2+^-binding

All-atom molecular dynamics simulations were performed using the GROMACS 5.1.2 package (Abraham et al., 2015). The simulation box contained one EH1.2 or EH1.2 Q382E molecule placed in a cubic box with a length of ~8 nm, which was filled with a 150 mM NaCl aqueous solution and which included additional Cl- ions to neutralize the whole system. The protein and ions were parameterized using the CHARMM36 force field (Huang and MacKerell, 2013). Newton’s equations of motion were integrated by employing the leap-frog algorithm (Hockney et al., 1974) with a time step of 2fs. The trajectory frames were recorded every 10ps. A cutoff of 1.2nm was applied to short-range electrostatic interactions while long-range electrostatics was calculated with the use of the particle mesh Ewald method (Darden et al., 1993). Van der Waals potentials were decreased so that the forces went smoothly to zero between 1.0 and 1.2nm. Bonds with hydrogen atoms were constrained by the LINCS algorithm (Hess et al., 1997) and water molecules were kept rigid by the SETTLE algorithm (Miyamoto and Kollman, 1992). The temperature of the system was maintained at 310K using the velocity rescaling thermostat with a stochastic term (Bussi et al., 2008) and the Parrinello–Rahman barostat (Parrinello and Rahman, 1981) was utilized for semi-isotropic pressure coupling with a reference pressure of 1.01bar. The time constants of the thermostat and barostat were 1ps and 5ps.

### TXRF

The EH domains were buffer exchanged using PD-10 columns to a buffer containing 20mM HEPES pH 7.4, 150mM NaCl, 0.5mM CaCl_2_. The proteins were concentrated to 2mM and the achieved protein concentration was verified by Nanodrop. TXRF quartz substrate disks were cleaned by placing them in a closed beaker with 5% HNO3 solution under boiling conditions for half an hour. This cleaning process was then repeated using a 3% HNO3 solution, rinsed twice using MilliQ H2O and a final rinse using a MilliQ H2O – ethanol solution and dried in vacuum. Five replicates for each EH domain consisting of 10 μL of the same protein solution were spotted on a quartz disk and dried under vacuum. The samples were measured in a G.N.R. TX2000 total reflection X-ray fluorescence spectrometer (40 kV, 30 mA, 1000 s LT, Mo anode). XRF data were fitted using the AXIL software package (Vekemans et al. 1994). Ca-Ka integrated intensities were normalized for the Cl-Ka integrated intensities to account for small fluctuations in X-ray tube current and amount of probed sample volume.

### Precipitation and refolding assays

To monitor precipitation and refolding by size exclusion chromatography, EH domains were treated with 10mM EGTA during 15 minutes, centrifuged at 10000g for 5min to remove precipitation and the soluble fraction was injected on a Superdex 75 10/300 (GE Healthcare). To refold EH domains, proteins were precipitated with 10mM EGTA and refolded overnight at room temperature by the addition of 50mM CaCl_2_. Proteins were injected on a Superdex 75 10/300 (GE Healthcare) to check the folded state of the protein.

To analyze the refolding of EH1.2 over time the protein was precipitated, in triplicate, using 10mM EDTA for 15 min while shaking followed by the addition of CaCl_2_ to a final concentration of 5 or 50mM after which the optical density at 600nm (Versamax, Molecular Devices) was measured over time.

### Liposome binding experiments

For the liposome binding experiments, a vesicle co-sedimentation assay was used as described in (Kooijman et al., 2007). The binding buffer was adapted to 20 mM HEPES, pH 7.4, 150 mM NaCl and a variable amount of CaCl_2_ was used.

### PolyPIPosome binding experiments

5μg protein was mixed with 40μl PolyPIPosomes (Echelon Biosciences) in a total volume of 80μl in a final buffer, constituted of 20mM HEPES pH 7.4, 150mM NaCl, 0.1mM CaCl_2_ and incubated for 1h at room temperature. After 1h, 30μl Monomeric Avidin Agarose beads (Pierce™) were added and incubated for 1h. The beads were spun down and washed three times with buffer. Liposomes were eluted from the beads by boiling at 95°C for five minutes after the addition of 10x sample reducing agent (Nupage, NP0009) and 4x Laemmli sample buffer (Biorad, #1610747).

### Live cell imaging and chemical treatments

Root epidermal cells of 5-7 day old seedlings, grown vertically on ½ MS medium in continuous light were imaged on a Leica SP8X microscope using the white light laser with a 40x/1.1NA water-immersion lens.

For sequential dual-color imaging, EGFP was visualized using 488nm laser excitation and a 495-550nm spectral detection. mRuby was visualised using 558nm laser excitation and a 600-700nm spectral detection. Time gating was always applied except for imaging FM4-64.

For PAO treatments seedlings were treated for 30min with 30μM PAO (Sigma, P3075, lot #MKCJ0095) in ½ MS media. For control experiments the same concentration of DMSO was used instead of PAO.

For ES9 treatment plants were pretreated for 5min with 10μM ES9, a generous gift of the lab of Jenny Russinova (PSB, VIB/Ugent, BE), in ½ MS media followed by 30min co-treatment of ES9 with 2μM FM4-64.

### PM/cytoplasm quantification

For the quantification of plasma membrane versus cytoplasm, the Fiji software package (v1.52p) was used. The 5% most intense pixelsof the ROI covering the PM were divided by the 5% most intense pixels of the ROI inside of the cell. Only images devoid of saturated pixels were used for quantification.

### Statistical analysis

For statistical analysis, the R package in R studio was used. Data were tested for normality and heteroscedasticity after which the multcomp package was used (Herberich et al., 2010).

### Peptidome profiling

Arabidopsis (Col-0) was grown vertically on ½ MS media for 7 days on a nylon mesh, harvested and flash-frozen in liquid nitrogen. Frozen seedlings (3g/experiment) were ground using mortar and pestle. Proteins were extracted and denatured in 50mM NaHCO3, 8M Urea and sonicated three times for 1min. Protein extracts were rotated for 30min at room temperature after which the extract was subsequently centrifuged twice at 20.000g for 20min. DTT was added to the supernatant, was added to a final concentration of 5mM and incubated at 55°C for 30min. Proteins were alkylated by the addition of 100mM iodoacetamide and incubated for 15min in the dark after which the mix was subsequently diluted with 50mM NaHCO3 to a final concentration of 2M urea. Trypsin was added in a ratio of 1:75, protein:trypsin (Sequencing Grade Modified Trypsin, V5117, Promega) and incubated overnight at 37°C on a rotating wheel.

Trypsin was removed using Sep-Pak Vac 3cc columns (500mg, WAT036815, Waters). The extract was acidified using 1% TFA for 15min on ice and cleared by centrifugation at 1780g for 15min at room temperature. The cleared extract was split into four and applied on an equilibrated Sep-Pak columns (Waters, WAT036815, Lot # 010437235B). The column was pre-wet using 5ml of 100% MeCN followed by sequential washing with 1 ml, 3 ml, and 6 ml of 0.1% TFA. After application of the extract, the column was washed sequentially with 1 ml, 5 ml, and 6 ml of 0.1% TFA followed by a 2ml wash with 0.1% TFA, 5% acetonitrile. The peptides were eluted using three times 2ml 0.1% TFA, 40% acetonitrile. The eluate was lyophilized for two days to remove TFA. 200μg EH domain or his-GFP were coupled, in triplicate, during one hour to 25μl Ni Sepharose 6 Fast Flow beads (GE Healthcare, 17-5318-01). Unbound protein was removed by three washing steps of 1ml each. The lyophilized peptides were solubilized in 20mM HEPES, 150mM NaCl buffer, 0.2mM CaCl_2_, divided and added to the coupled beads. The peptidome was incubated with the proteins for 4h after which the beads were washed three times with 1ml binding buffer. Peptides were eluted by the addition of 80μl 20mM HEPES pH 8, 8M Urea. The supernatant was removed from the beads and desalted with Monospin C18 columns (Agilent Technologies, A57003100) as described in (Leene et al., 2019).

Peptides were re-dissolved in 20 μl loading solvent A (0.1% TFA in water/ACN (98:2, v/v)) of which 5μl was injected for LC-MS/MS analysis on an Ultimate 3000 RSLC nano LC (Thermo Fisher Scientific, Bremen, Germany) in-line connected to a Q Exactive mass spectrometer (Thermo Fisher Scientific). The peptides were first loaded on *a* trapping column made in-house (100 μm internal diameter (I.D.) × 20 mm, 5 μm beads C18 Reprosil-HD, Dr. Maisch, Ammerbuch-Entringen, Germany) and after flushing from the trapping column the peptides were separated on a 50 cm μPAC™ column with C18-endcapped functionality (Pharmafluidics, Belgium) kept at a constant temperature of 35°C. Peptides were eluted by a linear gradient from 98% solvent A’ (0.1% formic acid in water) to 55% solvent B’ (0.1% formic acid in water/acetonitrile, 20/80 (v/v)) in 30min at a flow rate of 300 nL/min, followed by a 5min wash reaching 99% solvent B’. The mass spectrometer was operated in data-dependent, positive ionization mode, automatically switching between MS and MS/MS acquisition for the 5 most abundant peaks in a given MS spectrum. The source voltage was 2.2kV, and the capillary temperature was 250°C. One MS1 scan (m/z 400-2,000, AGC target 3 × 10^6^ ions, maximum ion injection time 80 ms), acquired at a resolution of 70,000 (at 200 m/z), was followed by up to 5 tandem MS scans (resolution 17,500 at 200 m/z) of the most intense ions fulfilling predefined selection criteria (AGC target 5 × 10^4^ ions, maximum ion injection time 80ms, isolation window 2Da, fixed first mass 140 m/z, spectrum data type: centroid, intensity threshold 1.3xE^4^, exclusion of unassigned, 1, 5-8, >8 positively charged precursors, peptide match preferred, exclude isotopes on, dynamic exclusion time 12s). The HCD collision energy was set to 25% Normalized Collision Energy and the polydimethylcyclosiloxane background ion at 445.120025Da was used for internal calibration (lock mass).

The raw files were processed with the MaxQuant software (version 1.6.4.0) (Cox and Mann, 2008), and searched with the built-in Andromeda search engine against the TAIR10_pep_20101214 database. Parameters can be found in the MS data file. Intensity values from the peptides output file of MaxQuant were used for quantitative analysis with the Perseus software (version 1.6.1.1). Intensity values were transformed to log2 values. Rows were filtered for at least 2 valid values in one of the sample groups, GFP, EH1.1 or EH1.2. Missing values were replaced with values from normal distribution with width of 0.3 and a downshift of 1.8. To determine the significantly enriched peptide sequences with the EH1 domains, a two-sided Student’s t test was performed between each of the EH domains versus GFP and the other EH domain as control. Permutation-based correction for multiple hypothesis testing was performed with thresholds FDR=0.01 and S0=1.

### NMR peptide binding

NPF peptide was dissolved in the final NMR buffer at a stock concentration of 4mM. ^1^H,^15^N HSQC titrations of ^15^N-labeled EH1.1 or EH1.2 with successive addition of unlabeled ligands were performed on samples containing 100 mM protein to a final concentration ratio of 1:4 excess of the peptide. Weighted chemical shift perturbations (CSP) were calculated as: 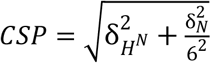 The CSPs were fitted to a binding isotherm using the equation:

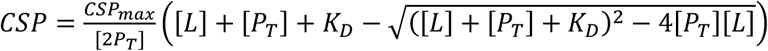

where CSP is the chemical shift perturbation at a given peptide concentration [L], CSPmax is the chemical shift perturbation at saturation, [PT] is the total protein concentration, and KD, the dissociation constant.

### BLI

BLI experiments were performed on an Octet RED96 instrument (FortéBio) using an HBS buffer supplemented with calcium (20mM HEPES pH 7.4, 150mM NaCl, 2mM CaCl_2_). A shake speed of 1000rpm at 25°C was used during all measurements. Ni-NTA (Molecular devices, 18-5102) biosensors were functionalized with EH-domains (10μg/ml) till a coupling signal of 3nm was reached. The proteins were covalently coupled to the Ni-NTA biosensors by sequentially dipping in a 20mM EDC:10mM NHS mix for 60 seconds followed by 60 seconds quenching in 1M ethanolamine pH 8.5. The coupled tips were equilibrated in buffer before the addition of the analyte. Next, functionalized sensors were sequentially dipped in increasing concentrations of analyte with an association time of 60s and a dissociation time of 120s. To correct for bulk effects during the measurements we performed double reference subtraction. Here, non-functionalized sensors were exposed to the same analyte concentrations while a functionalized sensor was dipped in zero concentration of analyte. The reference traces were subtracted from the raw data before analysis. Req values used in the analysis were determined for each concentration by averaging 30 data points once the sensors achieved stable equilibrium. Graphpad was used to analyse and plot binding data using the one-site total binding model. Peptides were ordered from Peptide 2.0 (95% purity).

### Visualisation of protein structures and data

For the visualisation of all protein structures UCSF Chimera was used. Mapping of conserved residues was performed by combining the Consurf server (Ashkenazy et al., 2016) with the generated alignment for the individual domains. All figures were prepared utilizing the Inkscape program (https://inkscape.org/).

### Accession codes

Coordinates and structure factors have been deposited in the Protein Data Bank under accession codes EH1.1 X-RAY PDB ID 6YIG

EH1.1 NMR PDB ID 6YEU, BMRB ID 34504

EH1.2 NMR PDB ID 6YET, BMRB ID 34503.

The final all-atom MD structure of EH1.2 and EH1.2 Q382E structure can be found in the Supplementary folder.

### Construction of transgenic plants

SCAMP5 and ΔN-SCAMP5 cDNAs with LR gataway sites were generated using a BioXP printer and cloned using LR clonase (Invitrogen) in pFASTRK-m43GW w/o terminator with a C-terminal GFP. Plant lines were generated by floral dip. Primary transformants were selected by fluorescent selection of the seeds and the seedlings were checked for expression level on a confocal microscope. T2 lines were used for subsequent experiments.

### Mutants and transgenic lines used in this study

Mutants and transgenic lines used in this study.
| Plant line | Background | Source |
| --- | --- | --- |
| UBQ10p::YFP-PH_FAPP1 (PI4P marker) | Col-0 | Simon et al. 2014 |
| 35Sp::AtEH1/Pan1-GFP | Col-0 | Gadeyne et al, 2014 |
| H3.3p::SCAMP5-GFP | H3.3p::AtEH1-mRuby in eh1/pan1(-/-) | This study |
| H3.3p::ΔN SCAMP5-GFP | Col-0 | This study |
| H3.3p::SCAMP5-GFP | Col-0 | This study |

### Primers

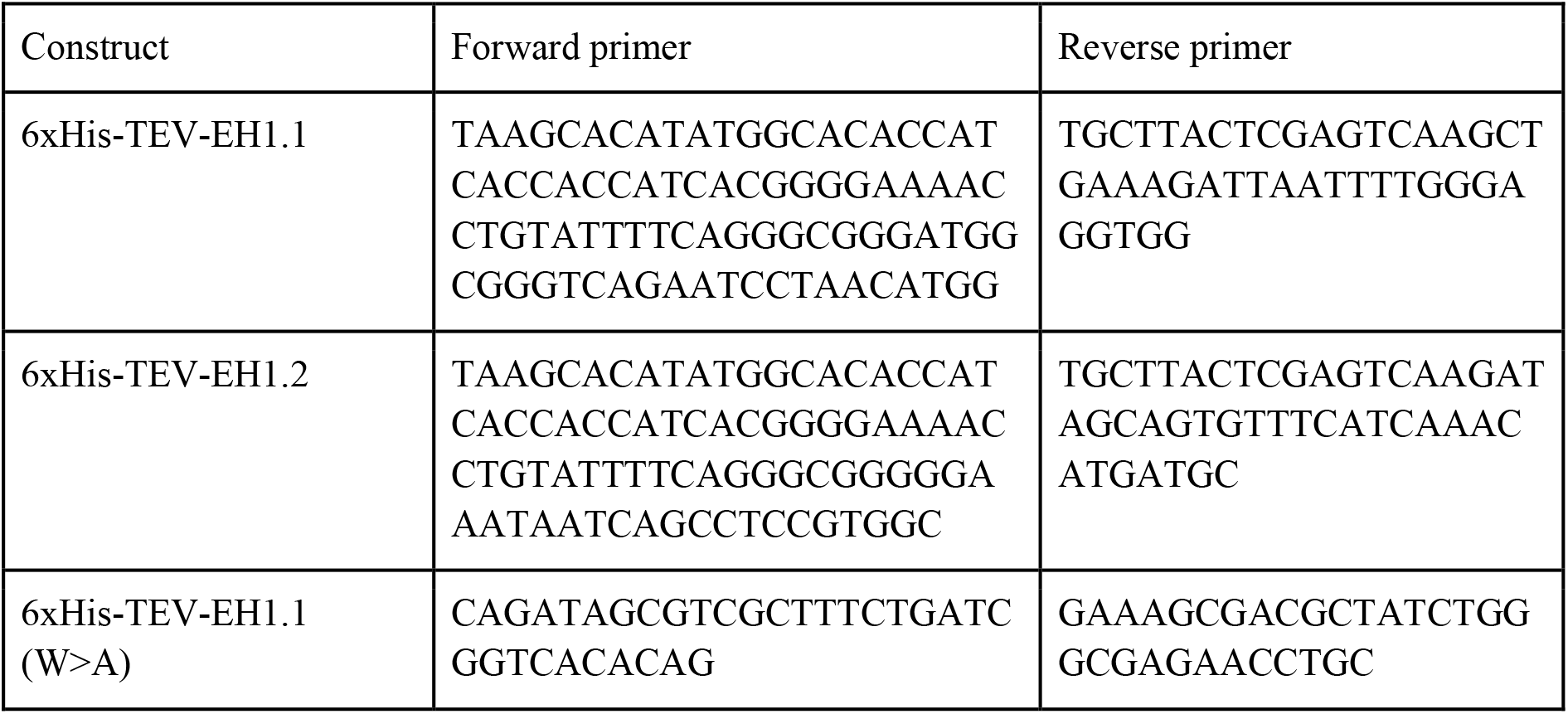

### BioXP generated sequences

> SCAMP5

~~~
TGTACAAAAAAGCAGGCTTAATGAATCGCCACCACGATCCCAATCCTTTCGATGAGGACGAAGAAATC
GTCAATCCTTTTTCGAAAGGTGGTGGAAGGGTTCCTGCTGCATCTAGGCCAGTTGAATATGGTCAAAG
CCTTGATGCTACTGTTGATATTCCATTGGATAATATGAATGACTCTTCACAGAAACAGAGAAAGCTTG
CTGACTGGGAAGCTGAGCTCAGGAAGAAAGAAATGGATATAAAGCGAAGAGAGGAAGCTATTGCTAAA
TTTGGTGTGCAGATAGATGATAAAAACTGGCCACCGTTTTTCCCAATCATACACCATGACATTGCTAA
AGAGATACCAGTTCATGCACAAAAGCTGCAGTATCTGGCTTTCGCTAGTTGGTTAGGTATCGTTCTGT
GTCTGGTATTCAATGTCATTGCAACGATGGTCTGCTGGATTAAAGGCGGAGGTGTTAAAATCTTTTTC
CTGGCCACAATATATGCATTGATCGGATGTCCACTCTCTTATGTACTATGGTACAGGCCACTCTACCG
AGCCATGAGGACTGACAGTGCTTTGAAGTTTGGTTGGTTTTTCTTCACCTACTTGATTCACATTGGCT
TCTGCATCGTTGCTGCCATCGCCCCTCCAATCTTTTTCCATGGAAAATCATTAACGGGTGTGCTTGCA
GCAATTGATGTCATCTCAGACAGTTTATTAGCTGGGATCTTCTACTTTATCGGATTCGGACTCTTCTG
CTTGGAGTCACTGCTGAGTCTATGGGTTCTTCAGAAAATTTACCTCTACTTTAGGGGAAACAAGTACC
CAGCTTTCTTGTACAA
~~~

> ΔN SCAMP5

~~~
TGTACAAAAAAGCAGGCTTAATGGGTGGTGGAAGGGTTCCTGCTGCATCTAGGCCAGTTGAATATGGT
CAAAGCCTTGATGCTACTGTTGATATTCCATTGGATAATATGAATGACTCTTCACAGAAACAGAGAAA
GCTTGCTGACTGGGAAGCTGAGCTCAGGAAGAAAGAAATGGATATAAAGCGAAGAGAGGAAGCTATTG
CTAAATTTGGTGTGCAGATAGATGATAAAAACTGGCCACCGTTTTTCCCAATCATACACCATGACATT
GCTAAAGAGATACCAGTTCATGCACAAAAGCTGCAGTATCTGGCTTTCGCTAGTTGGTTAGGTATCGT
TCTGTGTCTGGTATTCAATGTCATTGCAACGATGGTCTGCTGGATTAAAGGCGGAGGTGTTAAAATCT
TTTTCCTGGCCACAATATATGCATTGATCGGATGTCCACTCTCTTATGTACTATGGTACAGGCCACTC
TACCGAGCCATGAGGACTGACAGTGCTTTGAAGTTTGGTTGGTTTTTCTTCACCTACTTGATTCACAT
TGGCTTCTGCATCGTTGCTGCCATCGCCCCTCCAATCTTTTTCCATGGAAAATCATTAACGGGTGTGC
TTGCAGCAATTGATGTCATCTCAGACAGTTTATTAGCTGGGATCTTCTACTTTATCGGATTCGGACTC
TTCTGCTTGGAGTCACTGCTGAGTCTATGGGTTCTTCAGAAAATTTACCTCTACTTTAGGGGAAACAA
GTACCCAGCTTTCTTGTACAA
~~~

