## supplemental material for "Distinct EH domains of the endocytic TPLATE complex confer lipid and protein binding"

### SUPPLEMENTAL FIGURES

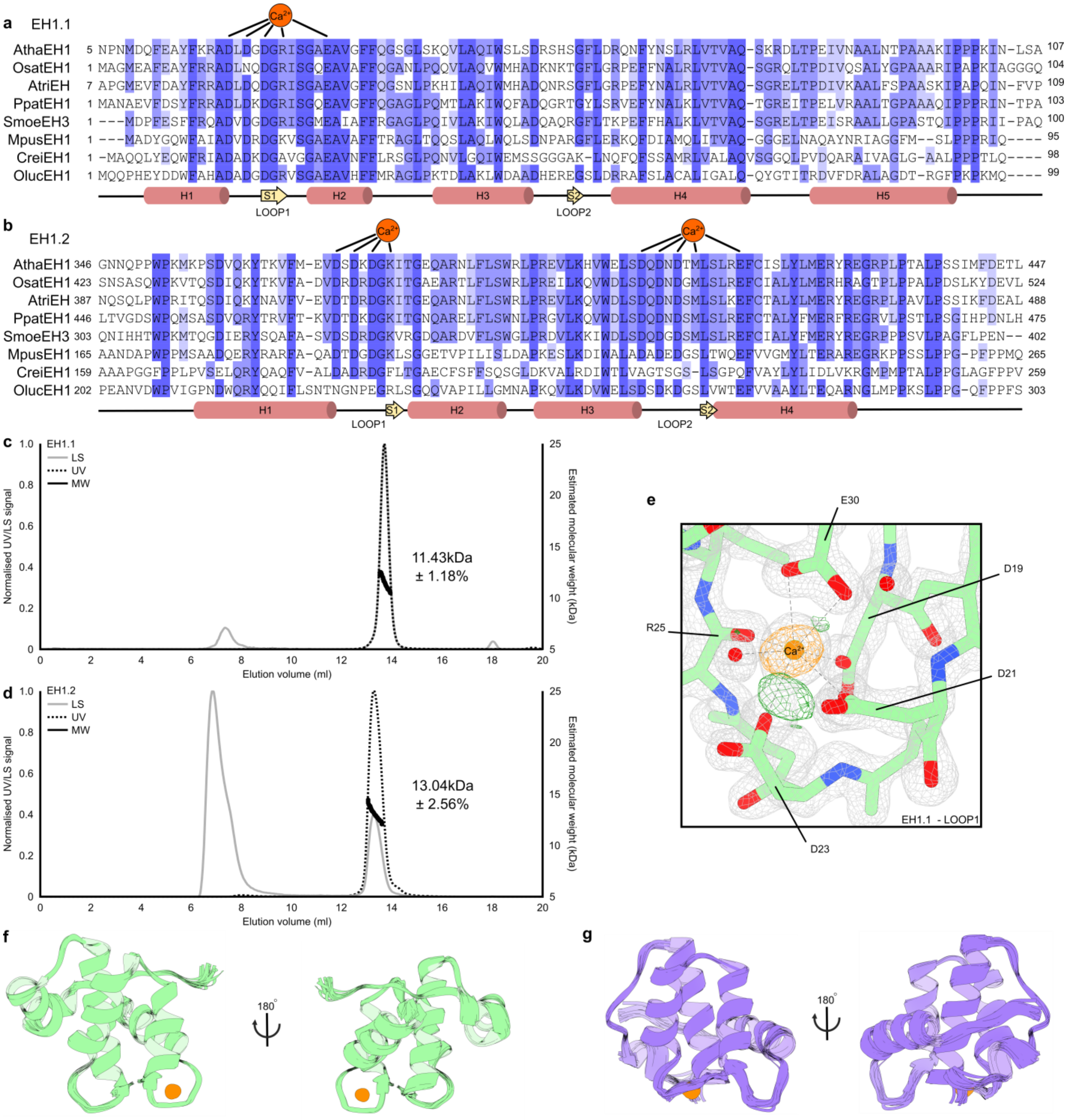

**Supplementary Figure 1: Both EH domains of AtEH1/Pan1 show $Ca^{2+}$ dependent reversible folding. a-b,** Detail of the MSA of the first and second EH domain of AtEH1/Pan1 (EH1.1 and EH1.2) of a selected number of species. $Ca^{2+}$ coordination is shown above and the secondary structural elements are indicated below each MSA. Atha - *Arabidopsis thaliana*, Osat - *Oryza sativa*, Atri - *Amborella trichocarpa*, Ppat - *Physcomitrella patens*, Smoe - *Selaginella moellendorffii*, Mpus - *Micromonas pusilla*, Crei - *Chlamydomonas reinhardtii*, Oluc - *Ostreococcus lucimarinus*. **c-d**, Size exclusion chromatography multi-angle laser light scattering elution profiles (Superdex 75 10/300), showing UV and light scattering signal (LS) of both EH domains of AtEH1/Pan1. The molecular weight distribution (MW) over the main peak is shown as black dots. **e**, Detail of the X-ray structure of the first EF-hand loop of EH1.1. Difference Fourier electron density map with coefficients 2mFo-DFc is shown as a grey mesh (contour level 2σ). Residual positive and negative difference electron density map with coefficients mFo-DFc (contour level

+-3σ) are shown in green and red (none present), respectively. Anomalous difference Fourier electron density for $Ca^{2+}$ is shown as an orange mesh (contour level 5σ). Side chains involved in $Ca^{2+}$ (orange sphere) coordination are showns as sticks. **f-g**, The ensemble of the 20 lowest energy NMR structures of EH1.1 (green, 5-106) and EH1.2 (purple,4351-448). Calcium is shown as a sphere (orange).

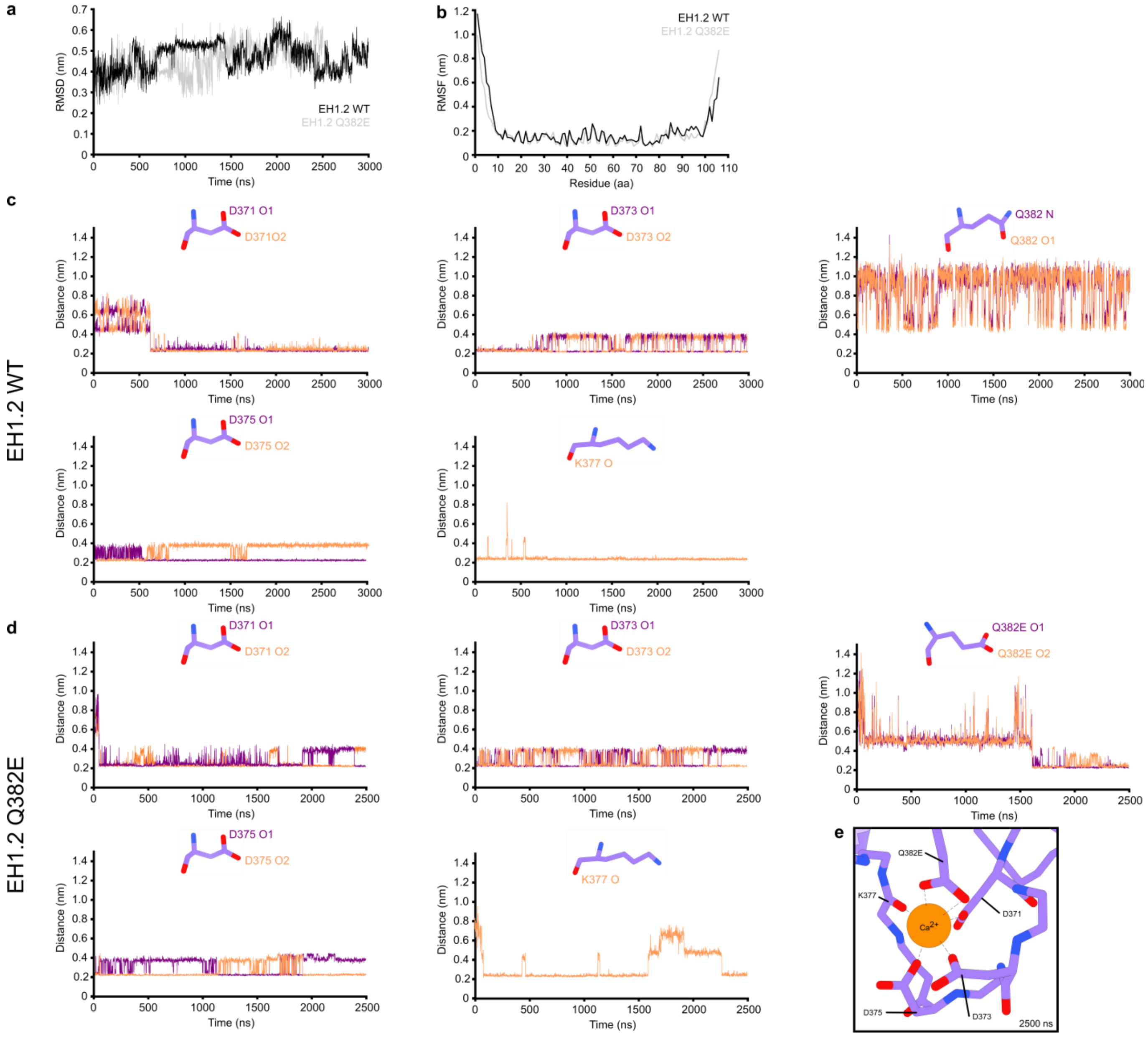

**Supplementary Figure 2: A possible coordination of the first loop of EH1.2 can be obtained by all-atom molecular dynamics simulations. a**, Root mean square deviation (RMSD) of EH1.2 WT (black) and the Q382E mutant (grey) over the simulation time. **b**, Root mean square fluctuation (RMSF) of EH1.2 WT amino acid residues (purple) and the Q382E mutant (orange) over the last 500ns of the simulation. The comparison of RMSF shows the flexibility of both protein termini for both WT and mutant structures. **c-d**, Oxygen to calcium (nitrogen and oxygen in the case of residue Q382) distance over the complete simulation of the residues potentially involved in $Ca^{2+}$-coordination. **e**, Cartoon representation of the first EF-hand loop of the all-atom MD structure of EH1.2 Q382E. $Ca^{2+}$ is shown in orange. The extensive molecular dynamics simulation did not reveal any contribution of Q382 in the coordination of calcium, in contrast to the simulation with the mutated EH1.2 domain, where canonical calcium coordination and hence contribution of Q382E was observed (defined as a 2.4 Å distance between calcium and the depicted atom).

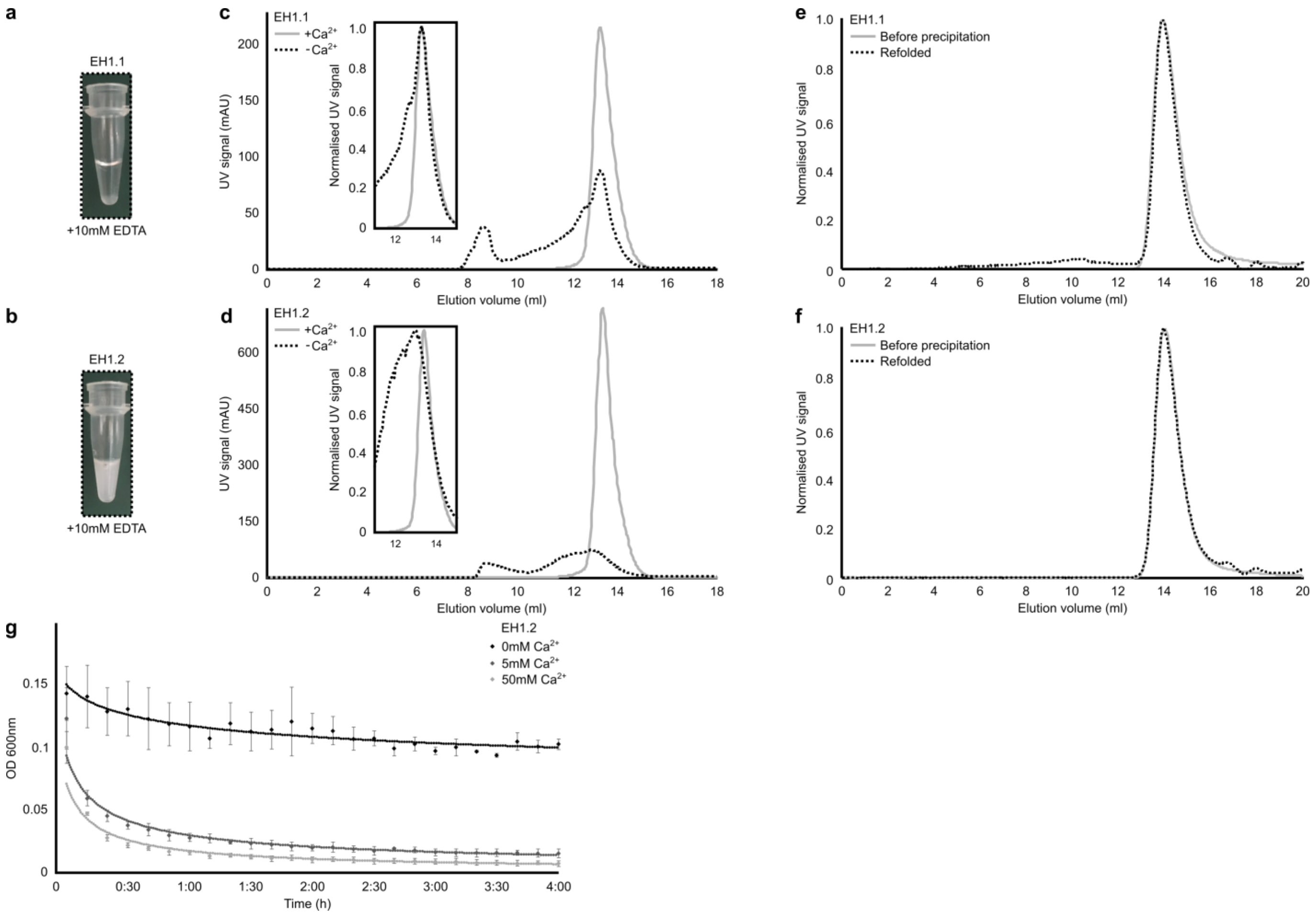

**Supplementary Figure 3: EH1.2 is more sensitive to calcium chelation than EH1.1. a-b**, Calcium chelation of EH domains. In contrast to EH1.1, $Ca^{2+}$ chelation (10mM EDTA) leads to extensive visible precipitation of EH1.2. **c-d**, Superimposed elution profiles (Superdex 75 10/300) of EH domains before and after $Ca^{2+}$ chelation (EGTA). Samples were centrifuged before injection to remove insoluble proteins. Normalized profiles are shown as inserts. Calcium chelation of EH1.2 results in complete precipitation compared to partial precipitation of EH1.1. **e-f**, Superimposition of elution profiles (Superdex 75 10/300) before precipitation and after refolding of both EH domains. EH domains were precipitated using 10mM EGTA and refolded. Elution profiles are normalized. **g**, Optical density measurement during refolding of EH1.2 in the presence of different $Ca^{2+}$ concentrations. The experiment was performed in triplicate, error bars show standard deviation.

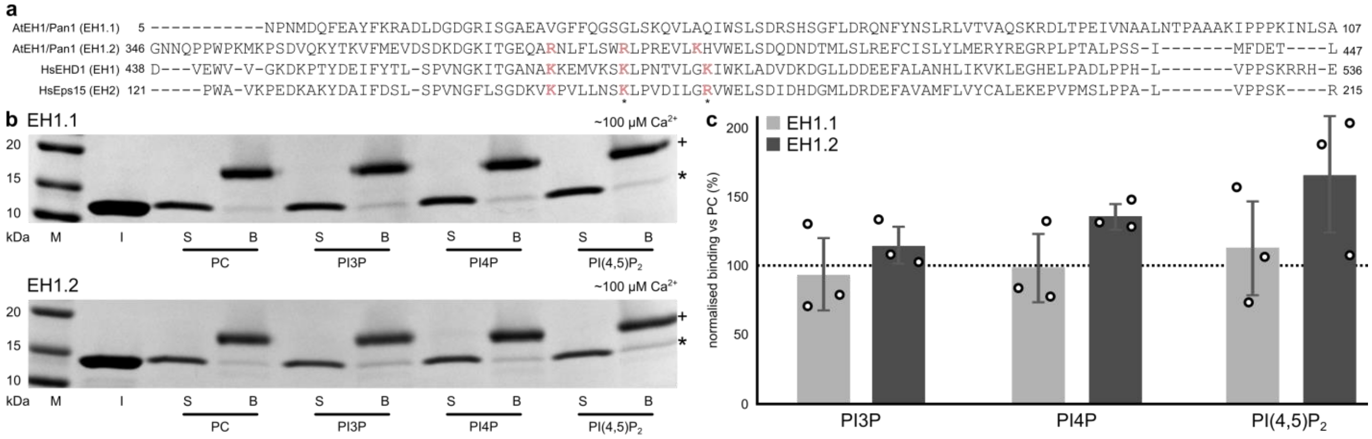

**Supplementary figure 4: Sequence alignment and PolyPiPosome binding of AtEH1/Pan1 EH domains. a**, Sequence alignment of the both EH domains AtEH1/Pan1 and EH domains shown to bind lipids by NMR (Naslavsky et al.). Conserved lysine and arginine residues are highlighted in pink. Residues shown to bind lipids by NMR are indicated with an asterisk (Naslavsky et al., 2007). **b**, PolyPIPosome binding of both EH domains of AtEH1/Pan1. HIS-EH domains were incubated with PolyPIPosomes and pelleted using mono-avidin beads. Bound (B) and supernatant (S) fractions were analyzed using Coomassie-blue stained SDS-PAGE gel (4-20%). Mono-avidin is indicated with plus (+) and the EH domain with an asterisk (*). Phosphatidylcholine (PC), Phosphatidyl-3-phosphate (PI3P), Phosphatidyl-4-phosphate (PI4P), Phosphatidyl-(4,5)-biphosphate (PI(4,5)$P_2$). M=Molecular marker, I=Input **c**, Quantification of data presented in panel b**,** (n =3). The ratio of supernatant (S) and bead (B) fractions was normalized to PC. Error bars show standard deviation.

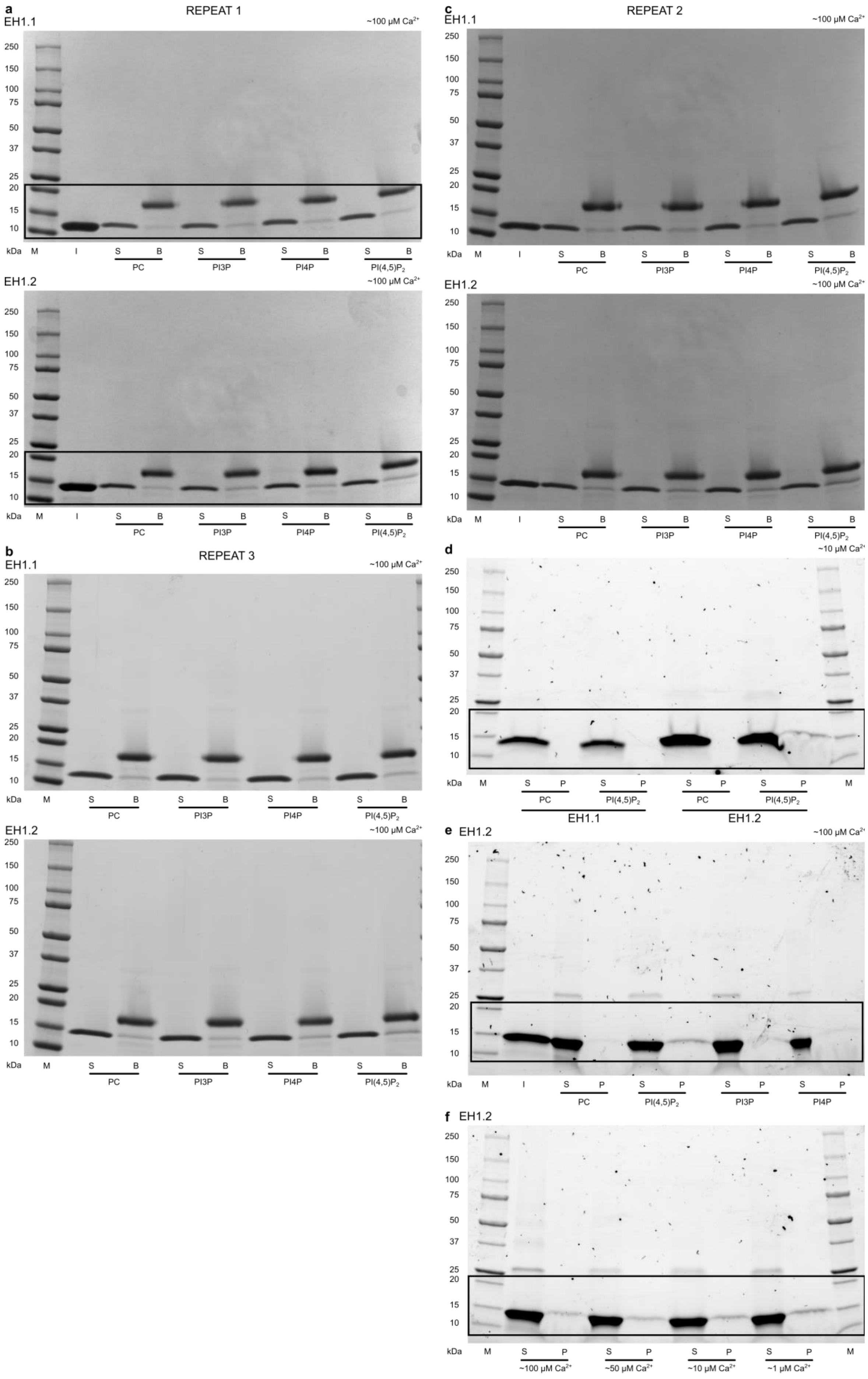

a
REPEAT 1
EH1.1
~100 µM Ca2+
250 150 100 75 50 37 25 20 15 10
kDa M I S B S B S B S B
PC PI3P PI4P PI(4,5)P2
EH1.2
~100 µM Ca2+
250 150 100 75 50 37 25 20 15 10
kDa M I S B S B S B S B
PC PI3P PI4P PI(4,5)P2
b
REPEAT 3
EH1.1
~100 µM Ca2+
250 150 100 75 50 37 25 20 15 10
kDa M S B S B S B S B
PC PI3P PI4P PI(4,5)P2
EH1.2
~100 µM Ca2+
250 150 100 75 50 37 25 20 15 10
kDa M S B S B S B S B
PC PI3P PI4P PI(4,5)P2
c
REPEAT 2
EH1.1
~100 µM Ca2+
250 150 100 75 50 37 25 20 15 10
kDa M I S B S B S B S B
PC PI3P PI4P PI(4,5)P2
EH1.2
~100 µM Ca2+
250 150 100 75 50 37 25 20 15 10
kDa M I S B S B S B S B
PC PI3P PI4P PI(4,5)P2
d
~10 µM Ca2+
250 150 100 75 50 37 25 20 15 10
kDa M S P S P S P S P M
PC PI(4,5)P2 PC PI(4,5)P2
EH1.1 EH1.2
e EH1.2
~100 µM Ca2+
250 150 100 75 50 37 25 20 15 10
kDa M I S P S P S P S P
PC PI(4,5)P2 PI3P PI4P
f EH1.2
250 150 100 75 50 37 25 20 15 10
kDa M S P S P S P S P M
~100 µM Ca2+ ~50 µM Ca2+ ~10 µM Ca2+ ~1 µM Ca2+

**Supplementary Figure 5: AtEH1/Pan1 lipid-binding depends on the second EH domain and is calcium-regulated. a-c**, Coomassie-blue stained SDS-PAGE (4-20%) analysis of PolyPiposome binding of both EH domains of AtEH1/Pan1. Uncropped blots of Supplemental Figure 4, panel b, as well as the other replicates of data quantified in Supplemental Figure 4, panel c. **d-f**, Expanded In-stain gel (4-20%) of liposome binding assays shown in Figure 2.

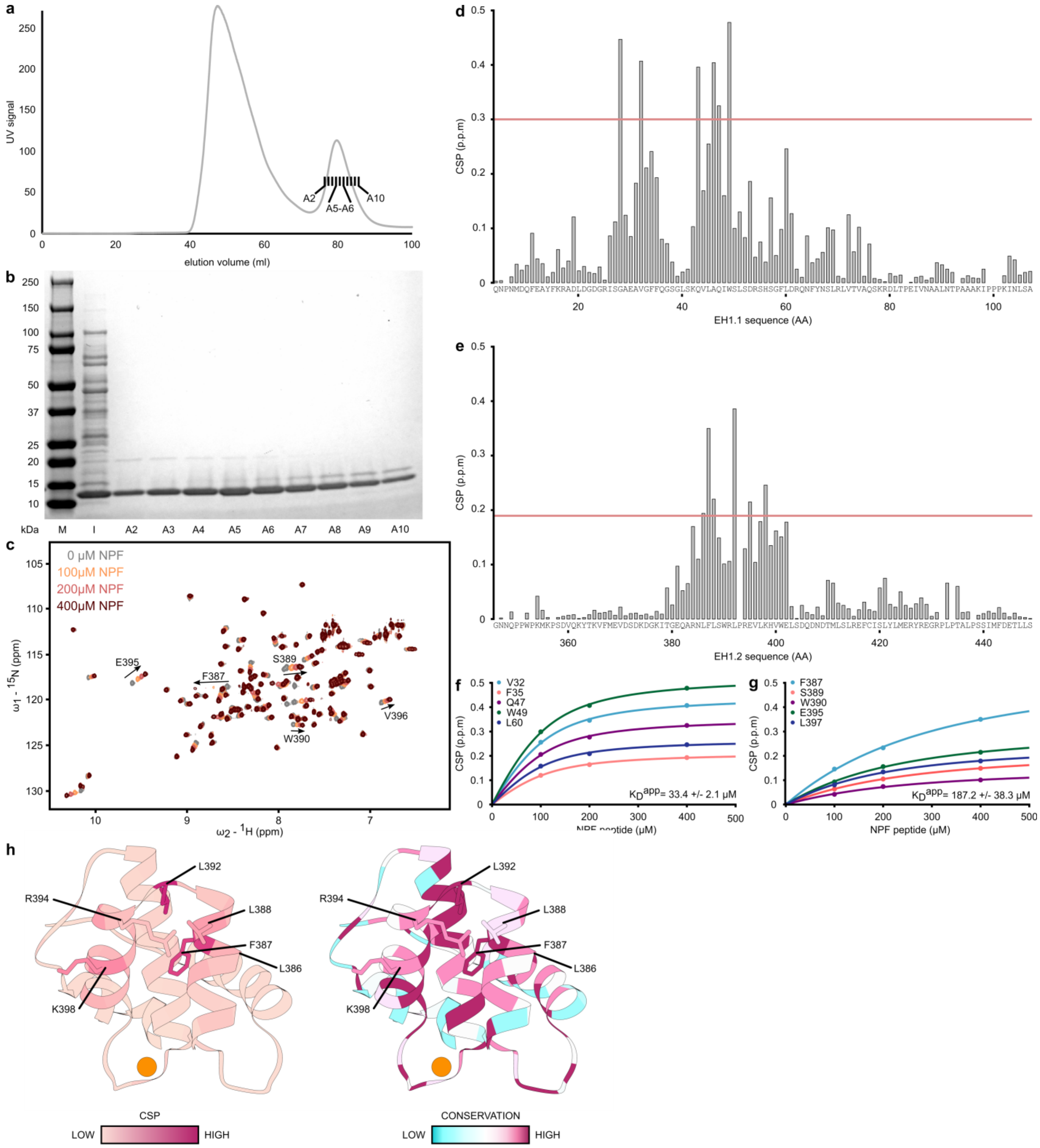

**Supplementary Figure 6: Elucidation of the interaction between SCAMP5 and AtEH1/Pan1.** **a**, Elution profile of EH1.1 W>A (Superdex 75 16/600 Hiload). **b**, Coomassie-blue SDS-PAGE (4-20%) analysis of the eluted fractions, shown in panel a, of the mutated EH1.1 domain (EH1.1 W49A) after size-exclusion chromatography (Superdex 75 16/600 HiLoad). **c**, $^{1}H$-$^{15}N$-HSQC spectra of EH1.2 showing chemical shift perturbations with increasing amounts of SCAMP5 NPF peptide (grey to red). The highlighted residues were used for Kd measurements in panel g. **d-e**, Chemical shift perturbation per residue of EH1.1 and EH1.2 upon the addition of 400μM double NPF peptide. The red line indicates the cut-off used in Figure 3, panel d-e. **f-g**, NMR binding analysis of the interaction between EH1.1/EH1.2 and the SCAMP5 N-terminal NPF peptide. The average dissociation constants ($K_D$) were estimated based on non-linear best fitting of the chemical shift perturbation (in ppm) of selected residues. **h**, Cartoon representation of the structure of the NMR obtained EH1.2 structure (351-448) colored according to chemical shift perturbations (in ppm) (left) or according to Consurf colors (right). Six residues showing the large chemical shift perturbations are shown as sticks.

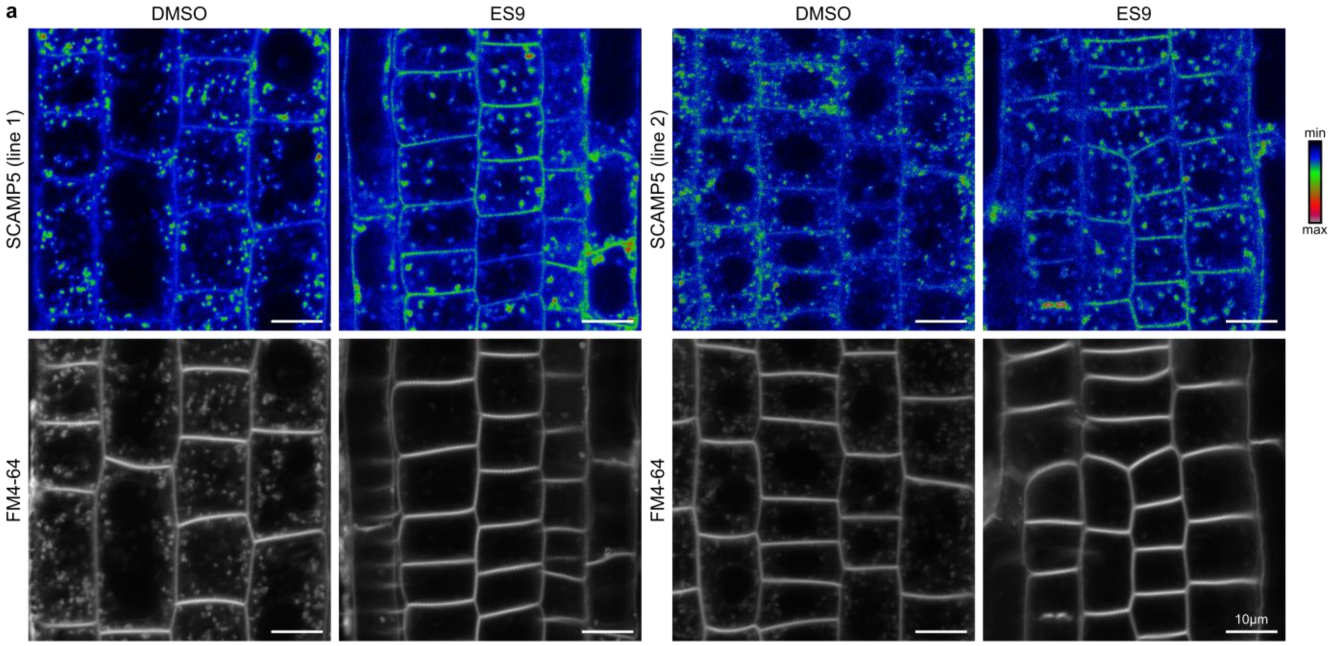

**Supplementary Figure 7: Disruption of endocytosis by ES-9 results in accumulation of SCAMP5 at the PM. a**, Short term ES9 treatment of two independent SCAMP5-GFP lines results in SCAMP5 PM accumulation. DMSO was used as a control. The styryl dye FM4-64 was used as a marker to monitor the effect of ES9 on the endocytic uptake. Scale bars indicates 10 µm.

#### SUPPLEMENTARY TABLES

**Supplementary Table 1: NMR and refinement statistics for EH1.1 and EH1.2. Structural statistics of 20 NMR models**

| | EH1.1 | EH1.2 |
|---|---|---|
| NMR distance and dihedral restraints | | |
| Distance restraints | | |
| Total NOE | 3127 | 2591 |
| Intra-residue | 570 | 588 |
| Inter-residue | 2557 | 2003 |
| Sequential ($\|i-j\| = 1$) | 732 | 615 |
| Medium-range ($1 < \|i-j\| < 5$) | 877 | 664 |
| Long-range ($\|i-j\| > = 5$) | 948 | 724 |
| Total dihedral angle restraints | 182 | 174 |
| φ | 91 | 87 |
| ψ | 91 | 87 |
| Structure statistics | | |
| Violations (mean ± s.d.) | | |
| Distance restraints (Å) | 0.021 ± 0.001 | 0.020 ± 0.001 |
| Dihedral angle restraints (º) | 0.243 ± 0.037 | 0.424 ± 0.046 |
| Max. dihedral angle violation (º) | 2.41 | 3.51 |
| Max. distance constraint violation (Å) | 0.363 | 0.322 |
| Deviations from idealized geometry | | |
| Bond lengths (Å) | 0.013 ± 0.000 | 0.012 ± 0.000 |
| Bond angles (º) | 1.43 ± 0.030 | 1.408 ± 0.030 |
| Impropers (º) | 1.43 ± 0.044 | 1.338 ± 0.044 |
| Average pairwise r.m.s. deviation* (Å) | | |
| Heavy | 0.63 ± 0.05 | 0.79 ± 0.07 |
| Backbone | 0.27 ± 0.04 | 0.41 ± 0.06 |
| Ramachandran plot statistics* (%) | | |
| Residues in most favoured regions | 92.4 | 94.4 |
| Residues in additionally allowed regions | 7.6 | 5.6 |
| Residues in generously allowed regions | 0.0 | 0.0 |
| Residues in disallowed regions | 0.0 | 0.0 |

* Residues 6-105 of EH1.1 and residues 351-448 of EH1.2

**Supplementary Table 2**: X-ray refinement statistics for EH1.1

| | EH1.1 |
|---|---|
| **Data collection** | |
| Beam line | Petra III-P14 |
| Wavelength (Å) | 1.033 |
| Space group | P $2_1$ $2_1$ $2_1$ |
| a,b,c (Å) | 35.51 38.62 63.52 |
| Resolution (Å) | 33 - 1.55 (1.605 - 1.55) |
| Total reflections | 177482 (7631) |
| Unique reflections | 12827 (1058) |
| Multiplicity | 13.8 (7.2) |
| Completeness (%) | 97.15 (83.02) |
| Mean I/σ(I) | 30.03 (4.69) |
| Wilson B-factor (Å$^2$) | 15.68 |
| R-merge | 0.0515 (0.3554) |
| R-meas | 0.05339 (0.3831) |
| R-pim | 0.01378 (0.1387) |
| CC1/2 (%) | 1 (0.962) |
| CC* | 1 (0.99) |
| **Refinement** | |
| Reflections in refinement | 12815 (1056) |
| Reflections used for R-free | 642 (53) |
| Rwork | 0.1449 (0.1944) |
| Rfree | 0.1640 (0.2462) |
| CC(work) | 0.968 (0.950) |
| CC(free) | 0.962 (0.973) |
| Number of non-hydrogen atoms | 889 |
| Number of macromolecular atoms | 792 |
| Ligands | 2 |
| Solvent | 95 |
| Protein residues | 198 |
| RMS bonds (Å) | 0.008 |
| RMS angles (Å) | 0.93 |
| Ramachandran favored (%) | 100 |
| Ramachandran allowed (%) | 0.00 |
| Ramachandran outliers (%) | 0.00 |
| Rotamer outliers (%) | 0.00 |
| Clashscore | 0.00 |
| Average B-factor (Å$^2$) | 22.05 |
| macromolecules | 20.70 |
| ligands | 19.56 |
| solvent | 33.32 |
| Number of TLS groups | 9 |

Statistics for the highest-resolution shell are shown in parentheses
Values reported by Phenix
Friedel pairs were treated as symmetry related
